# A statistical nonparametric method for identifying consistently important features across samples

**DOI:** 10.1101/833624

**Authors:** Natalie Sauerwald, Carl Kingsford

## Abstract

In many applications, a consistently high measurement across many samples can indicate particularly meaningful or useful information for quality control or biological interpretation. Identification of these strong features among many others can be challenging especially when the samples cannot be expected to have the same distribution or range of values. We present a general method called conserved feature discovery (CFD) for identifying features with consistently strong signals across multiple conditions or samples. Given any real-valued data, CFD requires no parameters, makes no assumptions on the shape of the underlying sample distributions, and is robust to differences across these distributions.We show that with high probability CFD identifies all true positives and no false positives under certain assumptions on the median and variance distributions of the feature measurements. Using simulated data, we show that CFD is tolerant to a small percentage of poor quality samples and robust to false positives. Applying CFD to RNA sequencing data from the Human Body Map project and GTEx, we identify housekeeping genes as highly expressed genes across tissue types and compare to housekeeping gene lists from previous methods. CFD is consistent between the Human Body Map and GTEx data sets, and identifies lists of genes enriched for basic cellular processes as expected. The framework can be easily adapted for many data types and desired feature properties.

**Availability:** Code for CFD and scripts to reproduce the figures and analysis in this work are available at https://github.com/Kingsford-Group/cfd.

**Supplementary information:** Supplementary data are available at https://github.com/Kingsford-Group/cfd.

## 1 Introduction

Many biological applications involve measuring features across a range of samples, raising the natural question of which features have consistently high measurements across these samples. Despite many methods for identifying features with significant differences between sets of samples [19, 18, 20], there is no general statistical method for identification of features that are consistent across sample sets. Similarity across conditions can signal the importance of a feature, such as an epigenetic mark required for proper gene regulation or an oncogene that is highly expressed across many cancer types. Genes that are highly expressed across many healthy tissue types are likely to be essential for cellular functioning, and can be used as controls for data normalization. This work presents a statistical method for identifying features that, across a majority of samples, have high measurements relative to their respective sample distributions.

Previous related methods focus on specific use cases, such as reproducibility measurements. Irreproducible discovery rate (IDR) is a statistical method to identify reproducible signals from high-throughput sequencing experiments, such as ChIP-seq peaks that are consistent between replicate experiments [13]. In principle this is a very similar goal to the one presented here, but the assumptions inherent to IDR are specific to the case of replicates. IDR looks for the top *n* signals with highest values, where the challenge is determining the *n* at which the correspondence between replicate signals drops off. When looking for similarity across non-replicate samples however, it is very likely that the set of consistent features will not be the most highly valued features in individual samples. Additionally, IDR expects the input samples to have similar distributions among the highest values, which is an appropriate assumption when looking at replicates but not when studying non-replicate samples.

Identification of features in single-cell data that are stable across cells has recently become an important problem as single-cell data becomes more available and prevalent in genomic and epigenomic analyses. There have been a few methods developed specifically for the discovery of so-called stably expressed genes (scSEGs) in single-cell RNA sequencing (scRNAseq) [9, 14]. Recently, a method called scMerge [14] used a Gamma-Gaussian mixture model to compute certain characteristics related to stability that are then used to rank genes by an “SEG index,” which is the average rank across these stability properties. The assumptions made by this model, including the Gamma and Gaussian priors, are specific to the analysis of scRNAseq and would likely not generalize to other domains. scMerge additionally requires an arbitrary rank percentile cutoff and assumes similar ranges of values across conditions. Another recent method called CORGI, which ranks genes based on their ability to capture common trajectories between scRNAseq data sets, is specifically for use in trajectory inference methods [24]. Though the goal of CORGI is to integrate data sets and therefore should be robust to distribution and value differences unlike scMerge, it optimizes for a different objective (capturing common trajectories) than the one stated here.

Genes that are consistently expressed across all cell and tissue types have been known as “housekeeping genes.” This term is generally used to describe genes that are required for basic cellular functioning, and many methods on many different data types have been developed for their identification [4, 7, 3, 21, 12, 26, 27, 5]. Housekeeping genes tend to be highly active, and their expression is essential for survival [25]. They can additionally be used as controls or in normalization methods for gene expression analyses. While they have been studied for a long time, there is little consensus on which genes are most confidently considered housekeeping genes, with differing methods reporting lists with little overlap [6, 21, 2]. One of the reasons for differing results is that there is no consistent definition of a housekeeping gene; some studies look for genes which are simply expressed in all samples [4], others look for genes with consistent expression levels across samples [7], and still others look for genes with high expression across samples [3]. The discrepancies between results from different studies highlights the importance of rigorous methods in this space.

## 2 Approach

We introduce a nonparametric statistical method, called conserved feature discovery (CFD), for identifying features with consistently high values across samples, robust to differences in distributions and ranges of values between the samples studied. CFD works with any data type that can be viewed as a list of features with numerical values for each sample. This includes RNA sequencing (RNA-seq) data, where features are genes or transcripts and the values represent their abundance, or ChIP-seq data, where features could be genomic bins and the values are the peak heights at these locations. In principle CFD could also be used with single-cell data, or with mass spectrometry measurements. Regardless of the underlying data type and feature set, CFD identifies the features that are statistically significantly conserved with relatively high values across input samples.

We provide theoretical guarantees on the specificity and sensitivity of CFD, and demonstrate its utility with simulation data to show tolerance to uninformative samples and false positives. We also apply CFD to two biological data sets, identifying housekeeping genes from human tissue RNA-seq data, resulting in robust gene lists enriched for annotations related to basic cellular processes.

## 3 Methods

Given a set of measurements on features in multiple samples, CFD identifies features that have consistently high values across samples by computing a conservation *p*-value that considers both consistency across samples and magnitude within a sample. The method first converts input values to rankings representing the fraction of the sample with higher values. The median and variance of these rankings for each feature are then combined with a *χ*^2^ test, and corrected for multiple hypothesis testing. CFD returns features that are statistically significant in conservation across samples and demonstrate relatively high values within each sample (Figure 1). Details of each step are given below.

**Figure 1:**
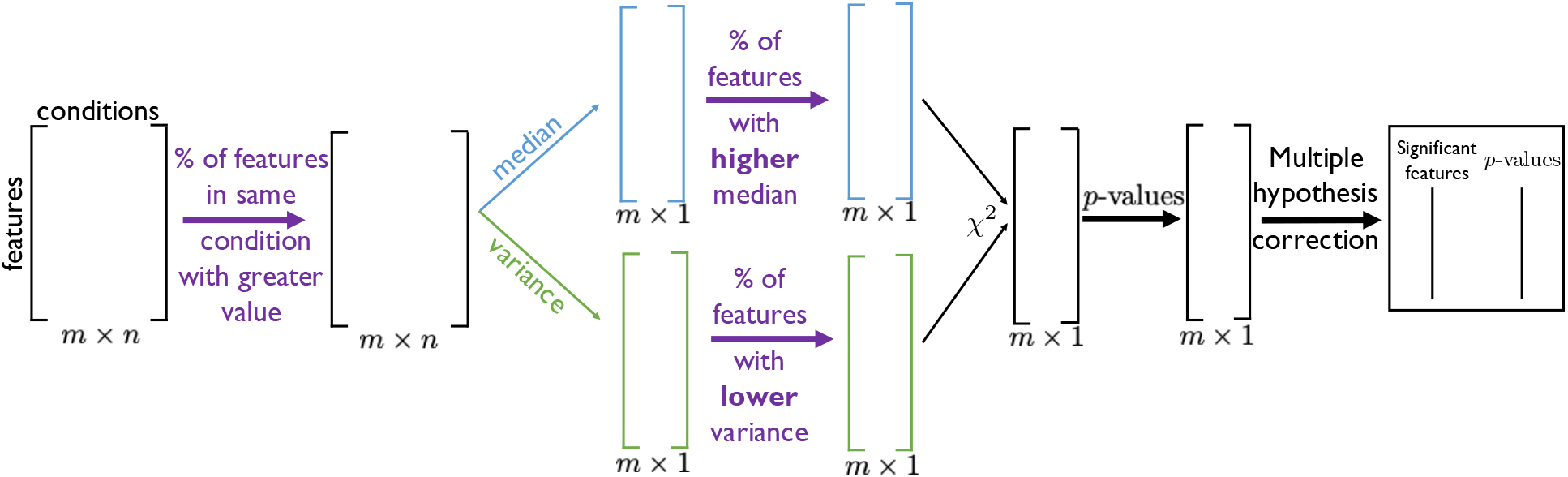
Method overview. The input to CFD is an *m* × *n* matrix of *m* feature measurements across *n* conditions. The given measurement values are converted to percentages based on their position within their respective sample distributions. Medians and variances of these values are taken across features, and converted to *p*-values which are combined with Fisher’s method to a *χ*^2^ value and converted back to *p*-values for multiple hypothesis correction. A list of statistically significant features and their *p*-values is returned.

### 3.1 Computing relative magnitude and variance of features

The input to CFD is a matrix **A** ∈ ℝ^*m*×*n*^ of measurements on *m* features, over *n* samples or conditions. We assume each sample has measurements drawn from some unknown underlying distribution that is well represented by the empirical data from that sample. We can therefore compute an empirical *p*-value without making assumptions on the underlying distributions of measurements. We define a function *ψ*:ℝ^*m*×1^ → [0, 1]^*m*×1^ that returns, for each value in the input vector, the fraction of the input vector with greater values, resulting in values between 0 and 1. We apply *ψ*(·) to each column of **A**, storing the results in a matrix **B** which is equal to the empirical cumulative distribution function subtracted from 1:

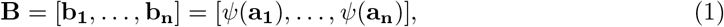

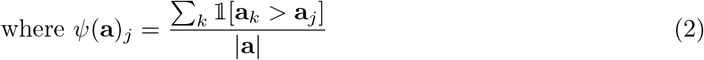

with 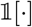 as the standard indicator function. This formulation makes CFD invariant to translations and positive multiplicative factors of the input samples. If the input data has negative values, it can therefore be shifted by any amount to ensure nonnegativity without altering the results. Given nonnegative data values, any features with zero values across all samples are removed to avoid testing unnecessary hypotheses where a feature takes the minimum of the domain under all conditions. This is particularly relevant in data such as RNA-seq, where many features (genes) will have zero values across all samples.

We compute the median and variance of each row of **B**. This gives two vectors **u** and **v** of length *m*, where **u**_*k*_ = median(**B**_*k*\*_) and **v**_*k*_ = Var(**B**_*k*\*_) for each row *k* of **B**. Intuitively, the vector **u** represents the median rank of each feature within its sample distribution, and **v** quantifies how much that ranking changes across samples.

### 3.2 Combination into one test statistic

We look for the features with high median ranking and low variance of rankings with respect to other features, computing **u**_*ρ*_ = *ψ*(**u**) and **v**_*ρ*_ = 1 − *ψ*(**v**), which measure how many features have higher median ranks (**u**_*ρ*_) and how many have lower variance of ranks (**v**_*ρ*_). The two vectors of independent probability values, taken as empirical *p*-values on the underlying distributions of each respective sample, **u**_*ρ*_ and **v**_*ρ*_, are combined with Fisher’s method, returning a vector **x** of *χ*^2^ values with four degrees of freedom:

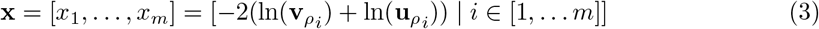

These values are then converted back to *p*-values for multiple hypothesis correction, via the standard cumulative density function for *χ*^2^ values.

### 3.3 Multiple hypothesis correction

At this stage, we have a vector of *p*-values that reflects the statistical significance of the medians and variances of all feature rankings within their sample distributions. However, many biological applications of this method will have a large number of features, requiring multiple hypothesis correction. We use the Benjamini-Hochberg procedure [1] to control the false discovery rate. Briefly, the Benjamini-Hochberg procedure returns an index on a list of sorted *p*-values, scaling the significance threshold based on the number of hypotheses tested (in this case, number of features), a parameter *α* for the false discovery rate, and the features’ position in the sorted list. Rather than adjusting the *p*-values themselves, this procedure results in a number of null hypotheses that can be rejected. We set *α* = 0.05, and report only the features and their *p*-values if the null hypothesis can be rejected.

## 4 Results

We first derive some theoretical guarantees on CFD’s sensitivity and specificity under certain assumptions, then demonstrate two desirable properties using simulated data. CFD is applied to biological data using two RNA-seq data sets, identifying genes that are consistently highly expressed across broad ranges of human tissue samples.

### 4.1 Theoretical guarantees

Assuming we are given features with median and variance ranks drawn from normal distributions, we show that with a probability dependent on the standard deviation of these distributions, the *p*-values of all true positive features will be less than 0.05 as long as the true positives make up no more than 10% of all features, and that all background features will have *p*-values greater than 0.05.

For the following, let there be *t* true positive features in a set *Y*, and *s* background features in a set *X*: *Y* ={*y*_*i*_ | *i* = 1, 2, …, *t*}, *X* = {*x*_*i*_ | *i* = 1, 2, …, *s*}. Suppose the background features have medians and variances drawn from normal distributions: 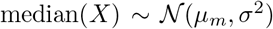, and 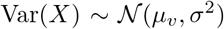. The true positives also have medians and variances drawn from normal distributions, with adjusted parameters. The medians of the true positives are assumed to be drawn from a normal distribution with a higher mean and lower standard deviation than the background: 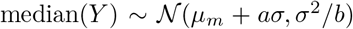. The variances of the true positives have lower mean and lower standard deviation than the background features: 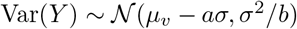. In particular, for simplicity in the following results we assumed that the true positives have medians drawn from a normal distribution centered two standard deviations higher than the background values with one third the standard deviation (*a* = 2, *b* = 3) ensuring that they will have generally higher values, and variances are drawn from a distribution centered at two standard deviations lower than the background, ensuring lower variance than the background. We will refer to the final *p*-value of a feature *x* computed by CFD as *p*_*cfd*_(*x*).

We first give a lemma that will be used later, defining the final *p*-value as a function of the median and variance ranks.

#### Lemma 1.

*For a feature ζ with p-value of its median rankings ϵ (there are ϵm features with a higher median) and p-value of its variance rankings δ (there are δm features with lower variance), the p-value p*_*cfd*_(*ζ*) *is given by p*_*cfd*_(*ζ*) = *ϵδ*(1 − ln(*ϵδ*)).

*Proof.* We first note that using Fisher’s method to combine *p*-values, the *χ*^2^ value for this feature will be *x* = −2(ln(*ϵ*) + ln(*δ*)) = −2 ln(*ϵδ*). In order to convert this value back to a single combined *p*-value *p*_*cfd*_(*ζ, x, k*), we must solve 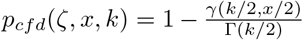, where *k* is the number of degrees of freedom (twice the number of *p*-values being combined) giving *k* = 4 here, Γ(·) is the standard Gamma function, and *γ*(·, ·) is the lower incomplete gamma function:

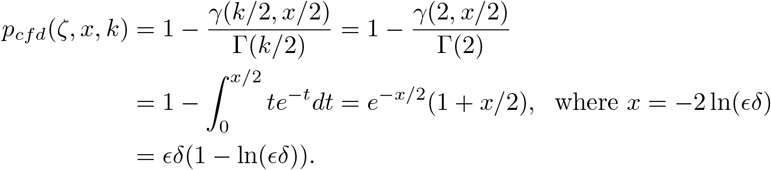

#### Proposition 1.

*With X and Y defined as above and* 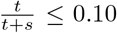, *p*_*cfd*_(*y*_*i*_) < 0.05 *for all true positive features y*_*i*_, *with probability at least* 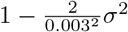.

*Proof.* For median values 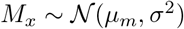 and 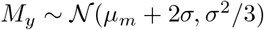,

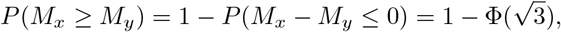

where Φ is the cumulative distribution function for the standard 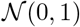 distribution. Similarly, for a variance values 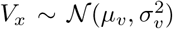 and 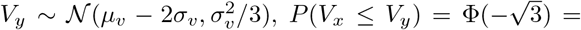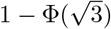.

For a particular true positive feature *y*, the probability of any other true positive feature having a greater median value than *y* is given by 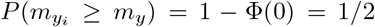 for any other *y*_*i*_ ∈ *Y*. The expected number of features with a higher median than *y* is therefore 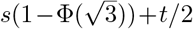, and by similar argument the expected number of features with lower variance is also 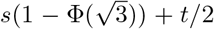. The total number of features is *s* + *t*, so both expected *p*-values of median and variance rankings for *y* are given by 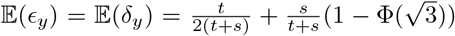. We note that *ϵ*_*y*_ = *δ*_*y*_ under our assumptions, so without loss of generality we will work with *ϵ*_*y*_ for the remainder of the proof. *ϵ*_*y*_ is a sum of two independent random variables (the fraction of background samples greater than *y* plus the fraction of other true positives greater than *y*), so its variance is given by: Var(*ϵ*_*y*_) = 2*σ*^2^/3 + 4*σ*^2^/3 = 2*σ*^2^. Using this fact and Chebyshev’s inequality, we can probabilistically bound the distance from *ϵ*_*y*_ to its expected value:

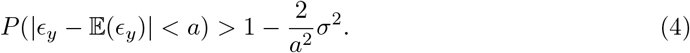

Rewriting the result of Lemma 1 as 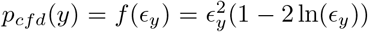, we now want to bound *p*_*cfd*_(*y*). For *ϵ*_*y*_ ∈ [0, 1], *f* (*ϵ*_*y*_) is Lipschitz with a constant of 1.5:

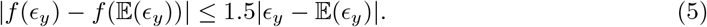

To bound the maximum *p*-value below 0.05, we therefore want to bound 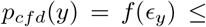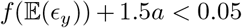. Setting *a* = 0.003 and using the assumption 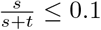, with probability at least 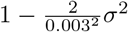:

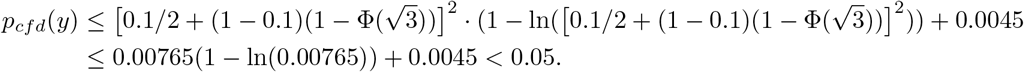

#### Proposition 2.

*With X and Y defined as above and assuming the existence of at least one true positive, p*_*cfd*_(*x*_*i*_) > 0.05 *for all background features x*_*i*_, *with probability at least* 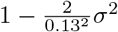.

*Proof.* For any background feature *x* with median *m*_*x*_, 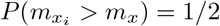 for any other *x*_*i*_, and 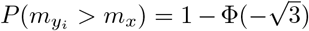 by analogous arguments used in the previous proof. The expected number of features with greater median, as well as those with lower variance, than *x* is therefore 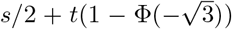. Using the same logic as above, 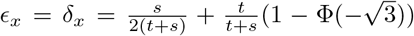. We again use Equation 4 and Equation 5, but now focus on the case where 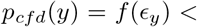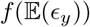, to bound *p*_*cfd*_(*y*) above 0.05.

Using the assumption that 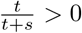 and setting *a* = 0.13,

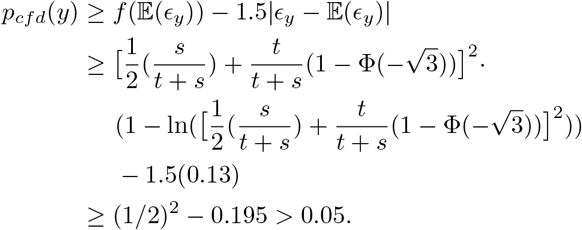

These bounds may not be practical because it is unlikely for biological data to be well represented by normal distributions, and in order for the probabilities we give to be high, the variance of the distributions (*σ*) must be extremely small. The Chebyshev inequality that these proofs depend on is quite weak, giving a loose bound, so more practical limits on the distributions may exist with a tighter bound. These bounds still provide some insight on the outcomes of CFD, notably showing that the conditions for avoiding false positives are much weaker than for guaranteeing discovery of all true positives.

### 4.2 Data

Simulated data was created to test two scenarios: tolerance of poor quality samples, and ability to avoid false positives. Matrices with 10000 features over 1000 samples were generated with 0%, 5%, 10%, 15%, 20%, and 25% of the samples drawn from a uniform distribution, which we will call uninformative samples. In each case, 50 features were drawn from a normal distribution with high mean and low variance (these are the true positives), and the rest of the features were drawn from normal distributions with other combinations of mean and variance values. For each percentage of uninformative samples, 100 matrices were simulated with true positives drawn from 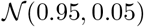. All other features were drawn from either 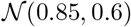 (high mean, high variance), 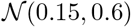 (low mean, high variance), or 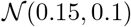 (low mean, low variance). Accuracy is measured as the true positive rate: the number of features correctly reported divided by the total number of true positive features.

For biological data, bulk RNA-seq from two studies was downloaded in .fastq format. The Illumina Human Body Map data consists of samples from 16 different normal human tissues, with two replicates of each. This is the same data used by Eisenberg and Levanon [7] to identify housekeeping genes previously. GTEx version 7 consists of 9781 samples from 55 different human tissue sites [15]. All bulk RNA-seq data was quantified as TPM values using Salmon version 0.9.1 [16]. We used the R package tximport [22] to combine transcript level quantifications to gene level quantifications, using gene annotations from GENCODE version 26 [11].

### 4.3 Performance on simulation data

On simulation data, CFD proved to tolerate a low percentage of poor quality samples, but its accuracy dropped to zero when 25% of samples were uniformly distributed. In all cases, CFD never reported any false positives so the specificity in each of these experiments was 100%. The fact that CFD does not achieve 100% accuracy on this simulation data even with 0% noisy samples is consistent with the observation from our theoretical results that identifying all true positives requires very strict assumptions on the data, which we did not satisfy in these simulations. While CFD maintains high accuracy for small proportions of noisy samples (there is no statistically significant difference in accuracy distributions between 0%, 5%, and 10% under the Kolmogorov-Smirnov two-sample test), the accuracy rapidly decreases when over 10% of samples do not preserve the high median and low variance of the true features, and goes to zero when 25% of samples do not match the conservation pattern (Figure 2). This result can also be interpreted to show that a feature cannot be considered “conserved” if its measurements are random in over 10% of samples.

**Figure 2:**
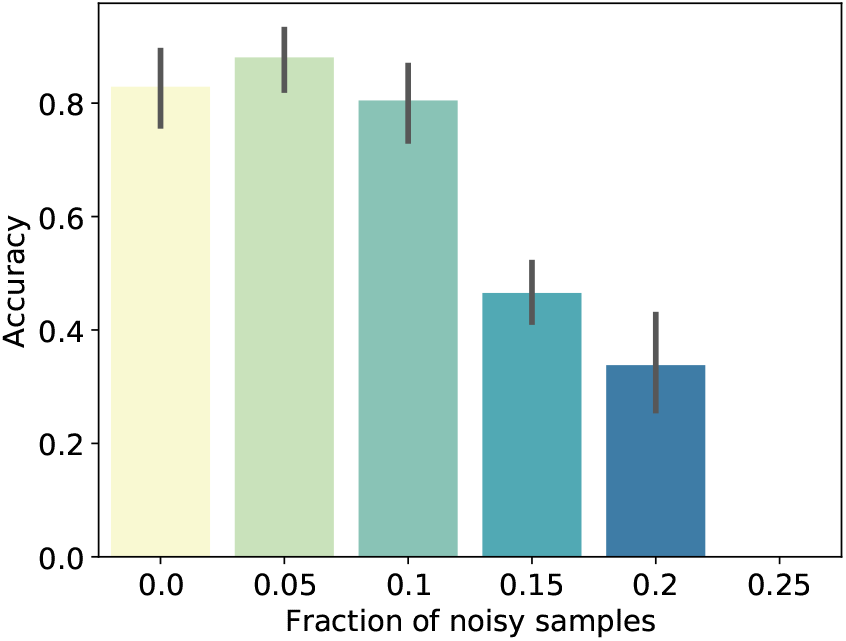
Performance on simulated data. Influence of the fraction of uniformly distributed samples on overall accuracy. Accuracy is measured as the percentage of true positives returned by CFD. Each bar represents the results from 100 simulated matrices, with the given percentage of samples drawn from a uniform distribution to simulate poor quality or uninformative samples. Error bars represent the standard deviation.

CFD is also able to identify cases in which none of the input fits the pattern of consistently high values. Previous methods such as scMerge [14] rank features by some conservation metric and pick the top *n*% as the conserved features, for some user-specified *n*. When no input features are truly consistently high, such an approach based on a parameter *n* will simply result in a list of *n* false positives. In contrast, we find that CFD returns no statistically significant features when all input features are either high mean and high variance, or low mean and low variance, across 14 different parameter settings (Table 1). Therefore in simulation data CFD appears highly robust to false positives, consistent with the theoretical guarantee on background features.

**Table 1:** Robustness to false positives. In each case, the given ratio of features was produced with high mean and high variance, with the remaining features drawn from distributions of low mean and low variance. Under all of these parameter settings, CFD returned no significant results.

| # feat. | # samp. | Hi mean | Low mean | Hi var | Low var | Ratio |
| --- | --- | --- | --- | --- | --- | --- |
| 10000 | 1000 | 0.7 | 0.3 | 0.4 | 0.2 | 0.5 |
| 10000 | 1000 | 0.7 | 0.3 | 0.5 | 0.1 | 0.5 |
| 10000 | 1000 | 0.85 | 0.15 | 0.5 | 0.1 | 0.5 |
| 10000 | 1000 | 0.85 | 0.15 | 0.4 | 0.2 | 0.5 |
| 20000 | 1000 | 0.7 | 0.3 | 0.4 | 0.2 | 0.5 |
| 20000 | 1000 | 0.7 | 0.3 | 0.5 | 0.1 | 0.5 |
| 20000 | 1000 | 0.85 | 0.15 | 0.4 | 0.2 | 0.5 |
| 20000 | 1000 | 0.85 | 0.15 | 0.5 | 0.1 | 0.5 |
| 20000 | 20 | 0.7 | 0.3 | 0.4 | 0.2 | 0.5 |
| 20000 | 20 | 0.7 | 0.3 | 0.5 | 0.1 | 0.5 |
| 20000 | 20 | 0.85 | 0.15 | 0.4 | 0.2 | 0.5 |
| 20000 | 20 | 0.85 | 0.15 | 0.5 | 0.1 | 0.5 |
| 10000 | 1000 | 0.85 | 0.15 | 0.5 | 0.1 | 0.25 |
| 10000 | 1000 | 0.85 | 0.15 | 0.5 | 0.1 | 0.75 |

### 4.4 Identifying housekeeping genes

CFD found a number of genes which are consistently highly expressed across human tissue samples on both the Human Body Map and GTEx data sets. On the Human Body Map data which consists of 16 samples, CFD ran in ≈1.7 seconds, and 8054 samples of GTEx data took ≈ 5 minutes on one core of a Linux Ubuntu 18.04 machine with an Intel Xeon Gold 6148 processor.

Housekeeping genes are typically defined as genes required for the maintenance of basic cellular functions, and they are expected to be relatively highly expressed in all cell and tissue types. Robust identification of these genes has proven challenging.

#### 4.4.1 Human Body Map

We identified 168 genes that passed statistical significance and multiple hypothesis testing using CFD. These genes are all consistently near the top of their respective sample distributions, demonstrating high values and low variance as desired. The top ten genes with lowest *p*-values are reported in Table 2, and the full list can be found in the Supplemental Data folder at https://github.com/Kingsford-Group/cfd. Nine of these genes are mitochondrially encoded genes. Mitochondrial DNA contains genes that are necessary for mitochondrial function [23], therefore it is reasonable to see these genes identified as consistently highly expressed across samples from various tissue types.

**Table 2:**
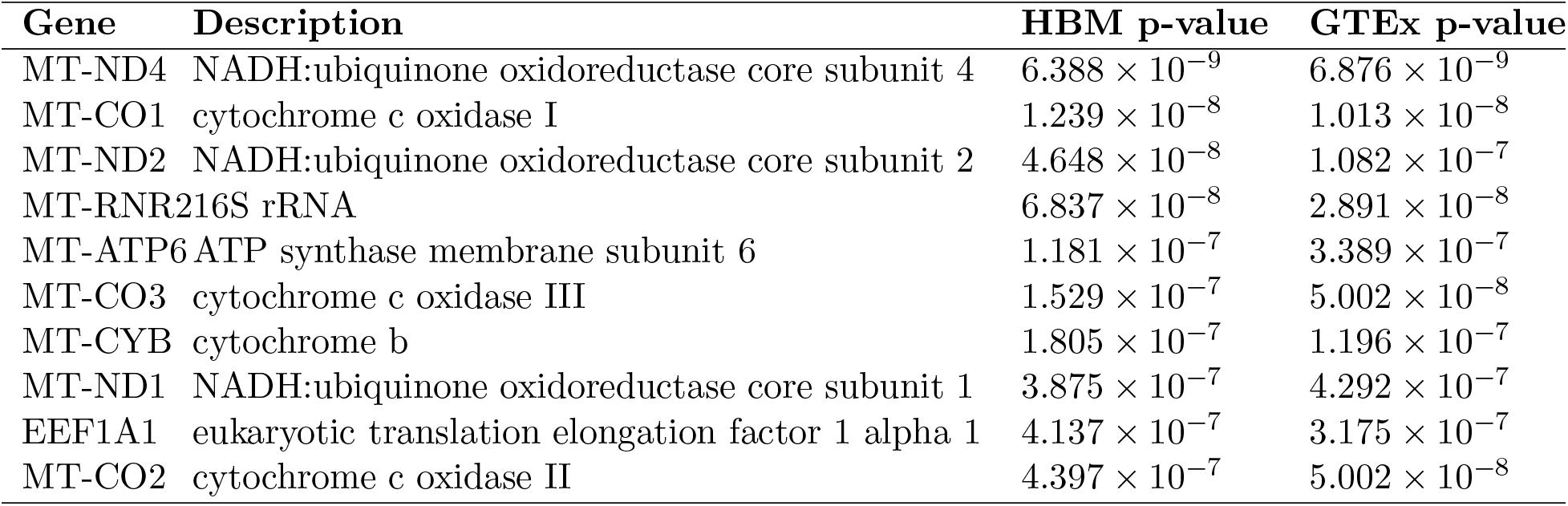
Top 10 most consistently highly expressed genes across human tissue samples computed by CFD, across the data from Human Body Map (HBM) and GTEx.

Our set of housekeeping genes is enriched for GO terms related to basic cellular functions and processes, across all three GO categories (Table 3). These results were obtained using gProfiler [17], with the ordered list of genes as input and the background as all human genes. Full GO enrichment results can be found in Supplemental Data at https://github.com/Kingsford-Group/cfd.

**Table 3:** Top three GO term results for each GO category on both Human Body Map and GTEx data.

| GO term | HBM $p_{adj}$ | GTEx $p_{adj}$ |
| --- | --- | --- |
| **Molecular Function** |  |  |
| RNA binding | $2.634 \times 10^{-53}$ | $2.010 \times 10^{-55}$ |
| structural constituent of ribosome | $2.851 \times 10^{-53}$ | $1.359 \times 10^{-66}$ |
| structural molecule activity | $2.033 \times 10^{-32}$ | $8.160 \times 10^{-42}$ |
| **Biological process** |  |  |
| SRP-dependent cotranslational protein targeting to membrane | $1.205 \times 10^{-66}$ | $1.525 \times 10^{-82}$ |
| translational initiation | $4.605 \times 10^{-66}$ | $1.931 \times 10^{-71}$ |
| cotranslational protein targeting to membrane | $2.656 \times 10^{-65}$ | $6.168 \times 10^{-81}$ |
| protein targeting to ER | $2.474 \times 10^{-63}$ | $1.359 \times 10^{-78}$ |
| **Cellular component** |  |  |
| cytosolic ribosome | $4.276 \times 10^{-66}$ | $1.825 \times 10^{-81}$ |
| ribonucleoprotein complex | $4.569 \times 10^{-54}$ | $6.926 \times 10^{-58}$ |
| ribosomal subunit | $2.896 \times 10^{-53}$ | $1.707 \times 10^{-66}$ |
| ribosome | $9.362 \times 10^{-53}$ | $7.568 \times 10^{-64}$ |

#### 4.4.2 GTEx: filtering out low quality samples

Not all of GTEx data is of high quality as noted in previous GTEx studies [10], so we filtered out lower quality samples in working with this dataset. With the help of MultiQC [8], we used mapping percentage (percent of reads that could be mapped during quantification) as a proxy for data quality and filtered GTEx by this value. Running CFD on the full, unfiltered GTEx version 7 release (9781 samples) returned no significantly conserved genes. Plotting the *p*-value distributions for the GTEx data shows that including the lower quality data, even filtering out samples with under 60% of reads mapped, produces an unexpected distribution with unexplained peaks (Figure 3**a**). This figure also suggests that many genes are very far from satisfying the desired property of high values across samples, as seen by the large number of genes with very high *p*-values. More significant genes could be obtained by filtering out some genes with *p*-values close to 1, thereby better satisfying the expectation of a uniform distribution of *p*-values and testing fewer unnecessary hypotheses. To verify that the genes we identified by thresholding the data were due to the higher data quality rather than simply a smaller sample size, we took ten random subsets of 8054 samples from GTEx, and ran CFD on each of them. None of these random subsets returned any significant genes, and all showed an uneven p-value distribution consistent with the full data (Figure 3**b**).

**Figure 3:**
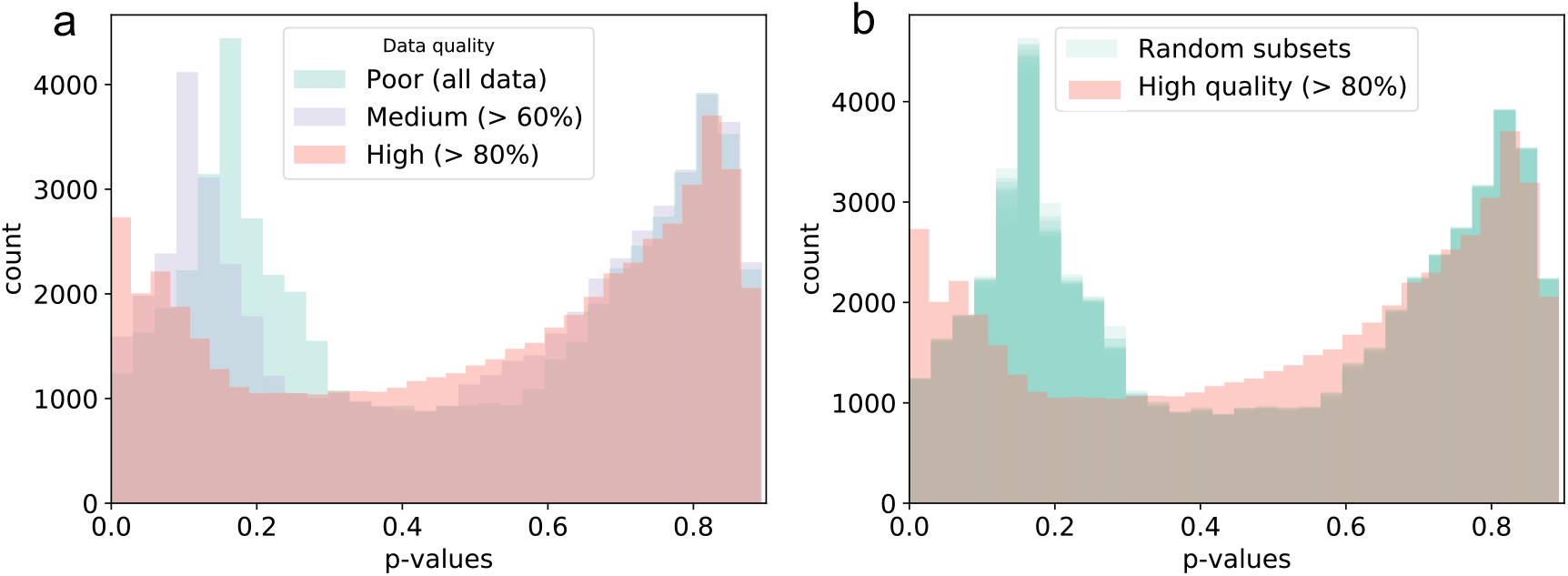
*P*-value histograms show the importance of filtering out low quality samples in GTEx data. Both plots show histograms of *p*-values across all genes as computed by CFD. **(a)** Comparison of *p*-value distributions from full GTEx data (9781 samples, green), partially filtered GTEx data (9216 samples, purple), and only the high quality samples (8054 samples, pink), as measured by mapping percentage. (b) Distributions from 10 random subsets of size 8054 from GTEx (green), as compared to the 8054 high quality (> 80% reads mapped) samples (pink).

On the subset of 8054 high quality GTEx samples, 149 genes passed statistical significance and multiple hypothesis testing. This gene list is enriched for similar terms as the set from the Human Body Map, again representing terms fundamental to cellular functioning. The top three terms from each category on each set (Human Body Map and GTEx) are provided with *p*-values in Table 3. For both the biological process and cellular component categories, the top three terms from Human Body Map and GTEx were not identical, so all terms in the top three of either list are reported. The top ten genes of the GTEx list are the same as the top ten we found from the Human Body Map data, though in a different order (Table 2).

### 4.5 Comparisons to previous work show little agreement in house-keeping gene lists

Housekeeping genes have been identified using many different data types and methods, with generally little agreement between them [6, 21, 2]. We compared our results with three studies from within the last ten years (Figure 4). Two of these previous studies [7, 4] identified many more housekeeping genes than we did (3804 and 2064 respectively), while the third resulted in a list of only 27 genes [3]. Chang et al. [4] computed housekeeping genes as those that are universally expressed in normal tissue, based on microarray samples. Eisenberg and Levanon [7] used the same Human Body Map RNA-seq data used here and defined house Chang et al. [4]keeping genes as those expressed at a constant level in all tissues. Caracausi et al. [3] searched for genes with high expression values and low standard deviation that were present in a large majority of samples, based on specific cutoff values for each criteria. These definitions all differ somewhat from each other and from the objective of CFD, which looks for genes that are consistently highly expressed across tissues, relative to their sample distributions. For both of our gene lists from GTEx and from the Human Body Map data, we find only 1 gene in common with all three previous lists (RPL8, a ribosomal protein), and about 50 genes in common with both Chang et al. [4] and Eisenberg and Levanon [7] (Figure 4**a**). Despite the different definitions, only approximately 30 genes in our set were not found in any of the previous lists, and half of these were mitochondrially encoded genes, which may not have been considered by previous studies.

**Figure 4:**
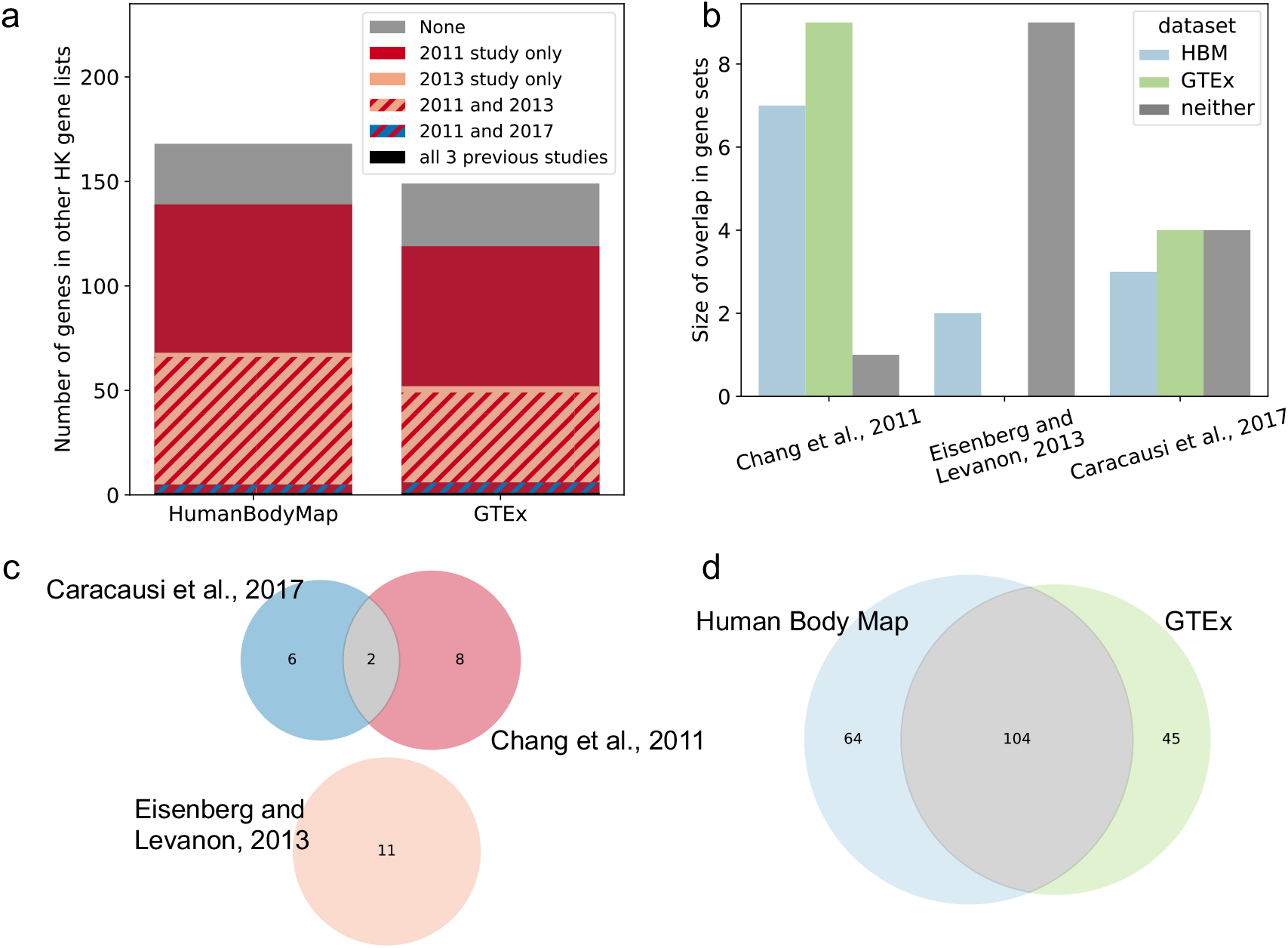
Comparisons with previous housekeeping gene lists. (a) Barplot showing the overlap between full HK gene lists from three previous studies (2011, 2013, and 2017 refer to Chang et al. [4], Eisenberg and Levanon [7], and Caracausi et al. [3], respectively) with our lists from Human Body Map and GTEx data. (b) Barplots showing the overlap with only the top 8-11 most confident genes reported by the same three previous studies. (c) Venn diagram of most confident gene sets from previous studies shows little agreement. (d) Venn diagram of the gene lists returned by CFD on Human Body Map and GTEx data shows significant overlap, suggesting our method is relatively consistent across these two input sets.

Most studies report a short list of their most confident housekeeping genes, and we see little consistency in these lists across methods. Caracausi et al. [3] gave a list of eight genes, and Eisenberg and Levanon [7] reported eleven. Chang et al. [4] did not provide a short list, but gave their full ranking, and we pulled the top ten from this list. Among the eight genes listed as best fitting the criteria of Caracausi et al. [3], four were not in either of the lists returned by CFD, despite the more similar definition of a housekeeping gene (Figure 4**b**). There is almost no overlap between the three previously published lists of “highly confident” housekeeping genes (Figure 4**c**); only two genes are shared between two studies, while the third study has no genes in common with either of the other two. Using CFD on our two data sets, we find a fairly large overlap in the full lists (Figure 4**d**), and as previously noted the top ten genes from both Human Body Map and GTEx are identical, suggesting that CFD returns fairly consistent results. Taken together, these results highlight the level of uncertainty and importance of methods in identifying housekeeping genes.

## 5 Discussion and Conclusions

We have introduced a general, nonparametric statistical method called CFD that identifies features with consistently high values across input samples, proved guarantees on its specificity and sensitivity, and demonstrated its effectiveness through simulated data and by identifying human housekeeping genes on two bulk RNA-seq data sets. CFD requires no parameters and makes no assumptions about the underlying distributions of or relationships between input samples, and has very high specificity. Simulated data suggests that CFD requires consistency across at least 80% of samples to identify any meaningful features. The housekeeping genes that we identify are consistent between two different RNA-seq data sets, and gene annotations suggest that the genes we identify are involved in fundamental cellular processes, as we would expect for housekeeping genes.

It is likely that more than the 149 or 168 genes identified by CFD satisfy the definition of housekeeping genes. The relatively low numbers reported here may be due to the very large number of genes that are either variable across tissue types or have low expression values, as shown by the large numbers of high *p*-values (Figure 3). If desired, more careful filtration of these genes that clearly do not fit the desired properties would likely yield more genes passing the multiple hypothesis testing procedure, and a larger list of housekeeping genes. The application to GTEx data in particular showed that CFD can sometimes benefit from some preprocessing or filtration steps to ensure the input data is not obscured by poor quality samples.

The framework of CFD can be easily adapted to identify any combination of high or low median and high or low variance features, simply by changing the direction of the inequality in *ψ*. In other applications, measurements with low values or high variance might be more of interest, and our statistical framework could be adapted in a straightforward manner to identify such features. In yet other applications it may be desirable to weight the relative importance of median and variance, which could be done by switching the *p*-value combination from Fisher’s method to Stouffer’s Z-score method, in which weights are straightforward to introduce.

This general statistical method represents a step towards principled analyses of conserved real-valued features across multiple conditions, and its framework can be easily adapted for different objectives. CFD could be applied to any data in which the same features are measured under different conditions, including gene expression, ChIP-seq, and protein quantifications.

## Acknowledgements

The authors would like to thank Jenn Williams for helpful discussions, as well as Hongyu Zheng for comments on the manuscript. The Genotype-Tissue Expression (GTEx) Project was supported by the Common Fund of the Office of the Director of the National Institutes of Health (commonfund.nih.gov/GTEx). Additional funds were provided by the NCI, NHGRI, NHLBI, NIDA, NIMH, and NINDS. Donors were enrolled at Biospecimen Source Sites funded by NCI Leidos Biomedical Research, Inc. subcontracts to the National Disease Research Interchange (10XS170), Roswell Park Cancer Institute (10XS171), and Science Care, Inc. (X10S172). The Laboratory, Data Analysis, and Coordinating Center (LDACC) was funded through a contract (HHSN268201000029C) to the The Broad Institute, Inc. Biorepository operations were funded through a Leidos Biomedical Research, Inc. subcontract to Van Andel Research Institute (10ST1035). Additional data repository and project management were provided by Leidos Biomedical Research, Inc.(HHSN261200800001E). The Brain Bank was supported supplements to University of Miami grant DA006227. Statistical Methods development grants were made to the University of Geneva (MH090941 & MH101814), the University of Chicago (MH090951,MH090937, MH101825, & MH101820), the University of North Carolina - Chapel Hill (MH090936), North Carolina State University (MH101819),Harvard University (MH090948), Stanford University (MH101782), Washington University (MH101810), and to the University of Pennsylvania (MH101822). The datasets used for the analyses described in this manuscript were obtained from dbGaP at http://www.ncbi.nlm.nih.gov/gap through dbGaP accession number phs000424.v7.p2.

